# Acquired resistance to the PRMT5 inhibitor confers collateral sensitivity to MEK inhibition in MTAP-null non-small cell lung cancer

**DOI:** 10.64898/2026.04.16.719008

**Authors:** Rongjie Fu, Yalong Wang, Ishita Rehman, Ella Bedford, Sana Sharif, Nghi D. Nguyen, Reid T. Powell, Andrew Adams, Weijun Liu, Shuyue Wang, Wei He, Yue Lu, Bin Liu, Pooja Anil Shah, Jordi Rodon Ahnert, Taiping Chen, Weiyi Peng, Clifford C. Stephan, Xinli Liu, Mark T. Bedford, Han Xu

## Abstract

Protein arginine methyltransferase 5 (PRMT5) is a synthetic lethal target in methylthioadenosine phosphorylase-deleted (MTAP-null) cancers. Second-generation MTA-cooperative PRMT5 inhibitors preferentially target MTAP-null cells while largely sparing MTAP-wildtype (MTAP-WT) cells, thereby improving tumor selectivity over first-generation PRMT5 inhibitors. Despite encouraging efficacy and safety signals in early clinical studies, the modest objective response rates (ORRs) observed with these inhibitors suggest that intrinsic or acquired resistance may limit their clinical benefit. Here, we investigated mechanisms of acquired resistance to the MTA-cooperative PRMT5 inhibitor BMS-986504/MRTX1719 in MTAP-null non-small cell lung cancer (NSCLC) cells and sought to identify therapeutic vulnerabilities that emerge upon resistance. Using multiple *in vitro*-derived resistant models, we found that acquired resistance was not fully explained by alterations in PRMT5 activity or reduced MTA levels. Instead, resistance was associated with collateral sensitivity to MEK inhibition and enrichment of MAPK-related transcriptional programs. Together, these findings identify MEK inhibition as an actionable collateral vulnerability in MTAP-null NSCLC cells that acquire resistance to PRMT5 inhibition.

## 1. Introduction

Homozygous deletion of chromosome 9p21.3 occurs in approximately 17% of human cancers (TCGA). Within this locus, the tumor suppressor cyclin-dependent kinase inhibitor 2A (*CDKN2A*) is the most frequently deleted gene (13.5%), followed by methylthioadenosine phosphorylase (*MTAP*) (9.3%). Because of their close genomic proximity, *MTAP* is commonly co-deleted with *CDKN2A*, with concurrent *MTAP*/*CDKN2A* loss observed in 9.2% of tumors across multiple cancer types, including glioblastoma, mesothelioma, urothelial carcinoma, pancreatic cancer, melanoma, and non-small cell lung cancer (NSCLC) (1,2). Loss of MTAP leads to accumulation of methylthioadenosine (MTA), which competes with the methyl donor S-adenosylmethionine (SAM) for binding to protein arginine methyltransferase 5 (PRMT5), thereby partially suppressing PRMT5 activity. In *MTAP*-deleted (MTAP-null) cancers, this reduction in basal PRMT5 activity creates a therapeutic window for further pharmacologic inhibition of PRMT5 relative to *MTAP*-wildtype (MTAP-WT) cells (3-8).

PRMT5 is a major type II protein arginine methyltransferase that catalyzes symmetric dimethylation (SDMA) of histone and non-histone substrates (9-11) and regulates diverse cellular processes, including transcription (12), alternative splicing (13), signal transduction (14), and the DNA damage response (15). PRMT5 is overexpressed in multiple cancer types, underscoring its therapeutic relevance (16,17).

First-generation PRMT5 inhibitors, including substrate-competitive and SAM-competitive compounds such as EPZ015666 (18) and JNJ-64619178 (19), exhibit limited selectivity between MTAP-WT and MTAP-null cancer cells because they effectively inhibit the SAM-bound form of PRMT5 that is also present in MTAP-WT cells. In contrast, second-generation MTA-cooperative PRMT5 inhibitors, such as BMS-986504/MRTX1719 (hereafter referred to as MRTX1719) (20-22), AMG 193 (23-25) and TNG908 (26,27), preferentially target the MTA-bound form of PRMT5 enriched in MTAP-null cancer cells while largely sparing MTAP-WT cells. This mechanism confers tumor-selective inhibition and helps overcome the limited therapeutic window associated with first-generation PRMT5 inhibitors. In phase I/Ib clinical trials, the tumor-selective PRMT5 inhibitors MRTX1719 and AMG 193 demonstrated objective response rates (ORRs) of 23% and 21.4%, respectively, with all reported responses being partial responses at well-tolerated dose levels (25,28). However, these modest ORRs suggest that intrinsic or acquired resistance may limit the clinical efficacy of this therapeutic strategy.

Acquired resistance to first-generation PRMT5 inhibitors has been described in multiple preclinical models, including EPZ015666-resistant murine and human NSCLC cells (29,30) and PRT-382-resistant mantle cell lymphoma (MCL) cells (31). Reported resistance mechanisms include upregulation of stathmin-2 (*STMN2*) or *PRMT5* mutations in EPZ015666-resistant cells, and activation of multiple signaling pathways, including mechanistic target of rapamycin (mTOR) signaling, in PRT-382-resistant MCL cells. Together, these findings highlight the need to define the molecular determinants of response to MTA-cooperative PRMT5 inhibitors and to identify rational therapeutic strategies that improve the depth and durability of clinical benefit.

In the present study, we developed *in vitro* models of acquired resistance to the MTA-cooperative PRMT5 inhibitor MRTX1719 in MTAP-null NSCLC cells. Using these models, we defined molecular features of the resistant state and identified candidate vulnerabilities arising during prolonged PRMT5 inhibition. Collectively, these findings provide preliminary insight into the molecular basis of acquired resistance and support further evaluation of targeting these vulnerabilities as a strategy to overcome resistance to MTA-cooperative PRMT5 inhibition in clinics.

## 2. Materials and Methods

### 2.1. Cell Culture

Human NSCLC cell lines H1299 (#CRL-5803, RRID: CVCL_0060, sex: male), H1975 (#CRL-5908, RRID: CVCL_1511, sex: female), A549 (#CCL-185, RRID: CVCL_0023, sex: male), H838 (#CRL-5844, RRID: CVCL_1594, sex: male), H1437 (#CRL-5872, RRID: CVCL_1472, sex: male), and H2126 (#CCL-256, RRID: CVCL_1532, sex: male); the human SCLC cell line H2171 (#CRL-5929, RRID: CVCL_1536, sex: male); human colorectal cancer cell lines RKO (#CRL-2577, RRID: CVCL_0504, sex: unspecified) and DLD1 (#CCL-221, RRID: CVCL_0248, sex: male); the human melanoma cell line A375 (#CRL-1619, RRID: CVCL_0132, sex: female); and the murine melanoma cancer cell lines B16-F10 (#CRL-6475, RRID: CVCL_0159, sex: male) were obtained from the American Type Culture Collection (ATCC). The murine fibrosarcoma cell line MCA205 (#SCC173, RRID: CVCL_VR90) and murine colon adenocarcinoma cell line MC38 (#SCC172, RRID: CVCL_B288, sex: female) were purchased from Sigma. All cell lines were maintained in the recommended media provided by the manufacturer at 37°C in a humidified incubator with 5% CO_2_ and were confirmed to be free of mycoplasma contamination prior to use using the MycoAlert Mycoplasma Detection Kit (Lonza, #LT07-705).

### 2.2. Drug Treatment

PRMT5 inhibitors (EPZ015666 and MRTX1719), MTA and MEK inhibitors (trametinib and selumetinib) were dissolved in DMSO. EPZ015666 was purchased from Sigma (#SML1421) and MRTX1719 from ChemiTek (#CT-MRTX1719). MTA was purchased from Sigma (#260585). Trametinib (#HY-10999) and selumetinib (#HY-50706) were purchased from MedChemExpress. Cells were treated with different concentrations of individual drugs or drug combinations for 6 days prior to western blot analysis and cell viability assays. DMSO was used as the vehicle control.

### 2.3. Generation of MTAP-isogenic Cell Lines

Human MTAP-knockout (MTAP-KO) cells were generated by infecting Cas9-expressing cells with lentiviruses produced from LentiGuide-Hygro-eGFP (Addgene, #99375) or LentiGuide-BSD vectors carrying sgRNAs targeting the human MTAP gene. Following hygromycin or blasticidin selection, single-cell clones were isolated, expanded, and validated for loss of MTAP expression by Western blot analysis. Murine MTAP-KO cells were generated by transfection of sgRNA:Cas9 ribonucleoprotein (RNP) complexes purchased from Integrated DNA Technologies (IDT) using Lipofectamine CRISPRMAX (Invitrogen, #CMAX00015). After transfection, single-cell clones were isolated, expanded, and validated for loss of MTAP expression by Western blot analysis. sgRNA sequences targeting human and mouse MTAP were designed using the GuidePro tool (32). The sgRNA sequences used were TCTGCCCGGGAGCTAAAACG for human *MTAP* and GGACAATAGTCACAATTGAG for mouse *MTAP*.

### 2.4. Generation of MRTX1719-resistant Cell Lines

1 x 10^5^ H838 cells, 1 x 10^5^ A549 cells and 2 x 10^5^ H1437 cells were seeded in 10-cm dishes and treated with MRTX1719 at initial concentrations of 0.02 μM for H838, 0.05 μM for A549, and 0.1 μM for H1437 cells. DMSO treatment was included for each cell line to control for potential solvent effects on drug response. Fresh drug-containing medium was replaced on Day 3 after treatment. On Day 7 post-treatment, cells were counted and re-seeded in 10-cm dishes at the same density as the initial seeding with the same drug concentration. After maintaining the cells at the same concentration for 4 weeks, the MRTX1719 dose was doubled and maintained for another 4 weeks. Using this dose-escalation strategy every 4 weeks, the final concentrations reached 1 μM for H838 and A549 cells and 2 μM for H1437 cells (6^th^ dose). The resulting MRTX1719-resistant (MRTXR) cells were subsequently maintained in drug-free medium and periodically tested for their response to MRTX1719 compared with DMSO-treated control cells using cell viability assays.

### 2.5. Cell Viability Assay and Synergy Scoring

To determine the optimal seeding density, human and murine cell lines were initially seeded in 96-well plates at different densities without drug treatment and monitored for 6 days using Incucyte (Sartorius) or Celigo (Revvity) confluence scanning. Based on the growth curves, cells were subsequently seeded in 96-well plates at the appropriate density in 100 µL complete medium and allowed to adhere for 14 hours prior to treatment. PBS was added to the outer wells of each 96-well plate to minimize edge effects.

The following day, each drug was serially diluted (3.162-fold dilution) in DMSO. Each dilution was supplemented with additional DMSO to maintain equal DMSO concentrations across all treatments. Cells were then treated with 100 µL of the single-drug dilution series and monitored daily using Incucyte. Each treatment condition was performed in triplicate. After 6 days of treatment, cell viability was measured using the CellTiter-Glo (CTG) assay (Promega, #G7571) according to the manufacturer’s instructions. Luminescence signals were recorded using a GloMax microplate reader (Promega). Percent inhibition values were calculated by normalizing the relative luminescence unit (RLU) values from treated wells to the average RLU values of DMSO-treated control wells. Dose-response curves were generated by plotting log(inhibitor) versus response using a variable slope (four-parameter) model in GraphPad Prism to determine IC50 values. For MTAP-isogenic cell pairs treated with EPZ015666 or MRTX1719, confluence measurements obtained from Incucyte scanning were used to calculate percent inhibition and IC50 values using the same normalization and curve-fitting methods as the CTG assay.

For drug combination treatments, MRTX1719 was combined with trametinib or selumetinib at the indicated concentrations. The concentrations were selected based on the responses observed in single-drug treatments. Cell viability was determined using CTG assay, and synergy mean scores were calculated using the Bliss independence model with the SynergyFinder+ tool (33). Scores < -10 indicate likely antagonistic interactions, scores between -10 and 10 indicate likely additive effects, and scores > 10 indicate likely synergistic interactions between the two drugs.

For MRTX1719 treatment with or without MTA supplementation, paired resistant cells were seeded in medium containing 10 μM MTA or no added MTA one day before MRTX1719 exposure. The MTA concentration was chosen based on the LC-MS analysis described below. On the next day, MRTX1719 was added as described above, and cell viability was measured using CTG after 6 days of treatment.

### 2.6. Western Blot

Cell pellets or adherent cells in culture plates were lysed using SDS lysis buffer (2% SDS, 10 mM Tris-HCl, 1 mM EDTA, pH 8.0), followed by heating at 98 °C for 10 minutes and vortexing every 2 minutes to ensure complete DNA shearing. Protein concentrations were determined using the Pierce BCA Protein Assay (Thermo Fisher Scientific, #23227) according to the manufacturer’s instructions. Equal amounts of protein were separated by SDS-PAGE and transferred onto nitrocellulose membranes using the Trans-Blot Turbo transfer system (Bio-Rad). Membranes were blocked in 5% milk in TBST and incubated with primary antibodies overnight at 4 °C on a rocking platform. The following day, membranes were incubated with HRP-conjugated secondary antibodies at room temperature for 1 hour. Chemiluminescent signals were detected using an Amersham ImageQuant 800 imaging system (Cytiva).

The following antibodies were used at the indicated dilutions: arginine methylation marks (SDMA, MMA, and ADMA; 1:2500) (34); PRMT5 (1:2000, Cell Signaling Technology, #79998, RRID: AB_2799945); MTAP (1:1000, Cell Signaling Technology, #4158, RRID: AB_1904054); phospho-MEK1/2 (Ser217/221) (1:1000, Cell Signaling Technology, #9154, RRID: AB_2138017); MEK1/2 (1:2000, Cell Signaling Technology, #9122, RRID: AB_823567); phospho-p44/42 MAPK (ERK1/2) (1:1000, Cell Signaling Technology, #9101, RRID: AB_331646); p44/42 MAPK (ERK1/2) (1:2000, Cell Signaling Technology, #4695, RRID: AB_390779); α-tubulin (1:5000, Developmental Studies Hybridoma Bank, #12G10); β-actin (1:5000, Sigma-Aldrich, #A1978, RRID: AB_476692); mouse IgG HRP-linked secondary antibody (1:10,000, Cytiva, #NA931, RRID: AB_772210); and rabbit IgG HRP-linked secondary antibody (1:10,000, Cytiva, #NA934, RRID: AB_772206).

### 2.7. Measurement of MTA levels by LC-MS

3 x 10^5^ H838 and 4.5 x 10^5^ H1437 resistant paired cells were seeded into individual wells of 6-well plates containing 1 mL of complete medium. Each cell line was seeded in biological triplicate. After 24 h, cell pellets and culture supernatants were collected for MTA quantification, and cell numbers were counted for normalization. MTA levels were measured using an AB SCIEX QTRAP 5500 mass spectrometer coupled to a Shimadzu UPLC system. Sample preparation and LC-MS/MS analysis were performed as previously described using MTA mass transition *m/z* 298.2 → 136.1 in positive ion mode (35). Intracellular (cell pellet) and extracellular (culture supernatant) MTA concentrations were quantified by comparison with a standard curve generated from external reference standards prepared in the mobile phase and normalized to the corresponding cell numbers.

### 2.8. High-throughput Drug Screening

A high-throughput drug screen was performed using a compound library consisting of 619 compounds, including 59 SGC epigenetic compounds, 380 TargetMol epigenetic inhibitors and 180 FDA-approved oncology drugs. Drug libraries were diluted in DMSO to generate 10 mM stock solutions and arrayed onto Echo-certified low dead volume (LDV) plates. The Combinatorial Drug Discovery Program (CDDP) at Texas A&M University maintains this compound library and performed the drug screening. For the screening assays, approximately 300 cells from H838 and H1437 resistant paired cell lines were seeded into Greiner black 384-well plates in growth medium using a Multidrop Combi liquid dispenser and cultured at 37 °C in a humidified incubator with 5% CO_2_ overnight. The next day, cells were treated with compounds by transferring materials from the LDV source plates into assay plates using the Labcyte Echo 550 platform. Cells were then exposed to the drug library at three concentrations (0.1 μM, 1 μM, and 10 μM) for 6 days. An 8-dose series of MRTX1719 was included as a positive control, and an 8-dose series of anisomycin was included as a nonselective control for drug response. At the endpoint, cells were washed, fixed, and stained with DAPI. Assay plates were imaged using a 4x objective on an ImageXpress microconfocal system, which captures the entire well area in a single field of view. Automated image analysis was performed using the advanced imaging collection in Biovia Pipeline Pilot to perform background correction and segment nuclei based on the DAPI signal. Drug screening was performed twice independently as biological replicates and drug responses between resistant paired cells were evaluated using two-way ANOVA analysis based on the Hafner growth rate index (GRI) (36) derived from cell counts to determine whether dose-response curves exhibited statistically significant drug effects. Significant drug candidates were identified based on response differences (MRTXR vs. DMSO) greater than 0.25 or less than -0.25 with an adjusted *p*-value < 0.05.

### 2.9. RNA sequencing and Data Analysis

RNA was isolated from H838 and H1437 resistant paired cells in triplicate using the RNeasy Mini Kit (QIAGEN, #74104) according to the manufacturer’s instructions. In-column DNase treatment was performed during RNA isolation. cDNA library preparation and RNA sequencing were performed by Signios Bio using the Illumina TruSeq Stranded mRNA Library Prep Kit, and sequencing was conducted on a NovaSeq X Plus platform. Each sample generated approximately 80 million paired-end reads of 150 bp in length. FASTQ files were mapped to the human reference genome (hg38) using TopHat (v2.0.10) and Bowtie (v2.1.0) with default parameter settings. Gene expression was quantified by transcripts per kilobase million (TPM) and transformed by log_2_ (TPM+1). Differentially expressed genes (DEGs) were identified by comparing MRTXR cells to DMSO cells using DESeq2 with thresholds of |log_2_ (Fold Change), FC| > 1 and a false discovery rate (FDR) < 0.05. KEGG pathway enrichment analysis was performed using the Metascape platform (37).

### 2.10. Statistical Analysis

Data are presented as mean ± SD. Statistical analyses were performed using GraphPad Prism (version 10). Statistical significance of changes in MTA levels was determined using an unpaired two-tailed t test. Statistical significance is indicated as follows: n.s., not significant; \**p* < 0.05; \*\**p* < 0.01; \*\*\**p* < 0.001.

## 3. Results

### 3.1 MRTX1719 selectively inhibits MTAP-null cells with variable responses

MRTX1719, an MTA-cooperative PRMT5 inhibitor, has been reported to selectively target MTAP-null cancer models and patient tumors (21). Consistent with this, MRTX1719 preferentially reduced viability in a panel of MTAP-null NSCLC cell lines after 6 days of treatment, whereas the substrate-competitive PRMT5 inhibitor EPZ015666 showed no MTAP-selective activity (**Figure 1A**). MTAP status and PRMT5 expression in these NSCLC models were confirmed by immunoblotting (**Figure S1A**).

**Figure 1.**
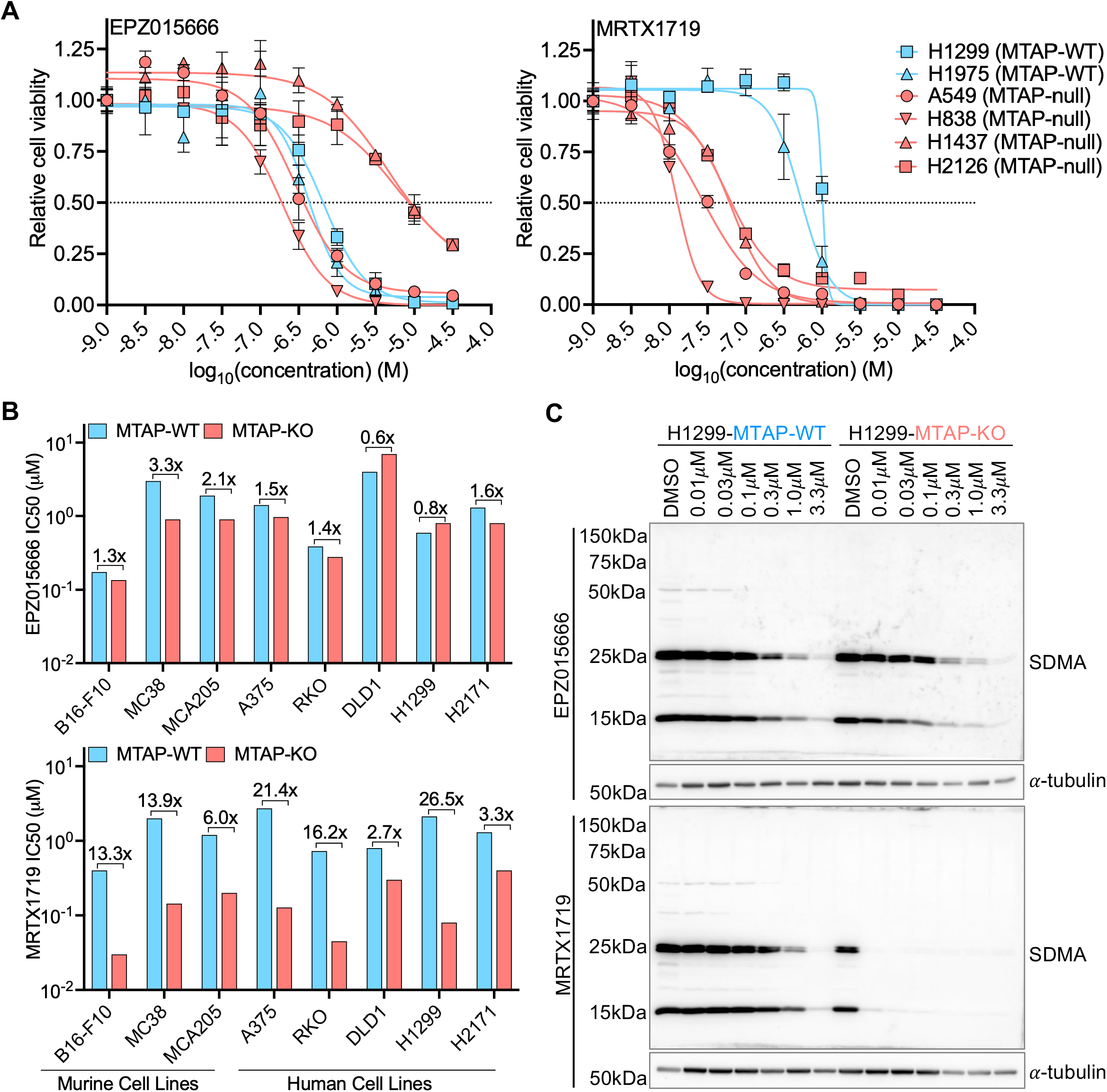
Selective but heterogeneous inhibition of MTAP-null cells by MRTX1719. **(A)** Dose-response curves of MTAP-WT and MTAP-null NSCLC cell lines treated with EPZ015666 or MRTX1719 for 6 days. Data are presented as mean ± SD. **(B)** Comparison of IC50 values for EPZ015666 (top) and MRTX1719 (bottom) in MTAP-isogenic murine and human cell lines. Fold differences in IC50 between MTAP-WT and MTAP-null cells are indicated. **(C)** Immunoblot analysis of SDMA levels in the H1299 MTAP-isogenic cell pair following treatment with increasing concentrations of EPZ015666 or MRTX1719.

To directly assess the contribution of MTAP loss to drug selectivity and sensitivity, we generated eight murine and human MTAP-isogenic cell line pairs spanning multiple tumor types using CRISPR-mediated MTAP knockout and confirmed MTAP loss by immunoblotting (**Figure S1B, Supplementary Table 1**). Following 6 days of treatment, MRTX1719 generally produced a greater IC50 shift (2.7-fold to 26.5-fold) between MTAP-WT and MTAP-KO cells than EPZ015666 (0.6-fold to 3.3-fold), although the magnitude of this effect varied substantially by model (**Figure 1B**). Notably, human DLD1 and H2171 MTAP-KO cells showed relatively limited sensitivity to MRTX1719 compared with other MTAP-KO models, suggesting that MTAP loss alone does not uniformly predict sensitivity in MTAP-null contexts.

Consistent with these growth effects, MRTX1719, but not EPZ015666, selectively reduced PRMT5 activity, as reflected by decreased SDMA levels, in H1299 and A375 MTAP-KO cells (**Figure 1C, S1C**). By contrast, MTAP-WT cells were largely unaffected by MRTX1719 at concentrations below 1 μM. Together, these findings establish that MRTX1719 preferentially inhibits cell viability and PRMT5 activity in MTAP-null cells, although sensitivity varies across models.

### 3.2 Prolonged MRTX1719 treatment induces acquired resistance in sensitive NSCLC cells

To investigate the mechanisms underlying MRTX1719 resistance, we used a dose-escalation strategy (38,39) to generate resistant cells *in vitro*. Sensitive NSCLC cell lines A549, H838, and H1437 were initially treated with MRTX1719 at approximately their viability IC50, and the dose was doubled every 4 weeks until reaching 1 μM in A549 and H838 cells and 2 μM in H1437 cells over 25 weeks (**Figure 2A**). Parallel long-term DMSO treatment served as a control for prolonged solvent exposure.

**Figure 2.**
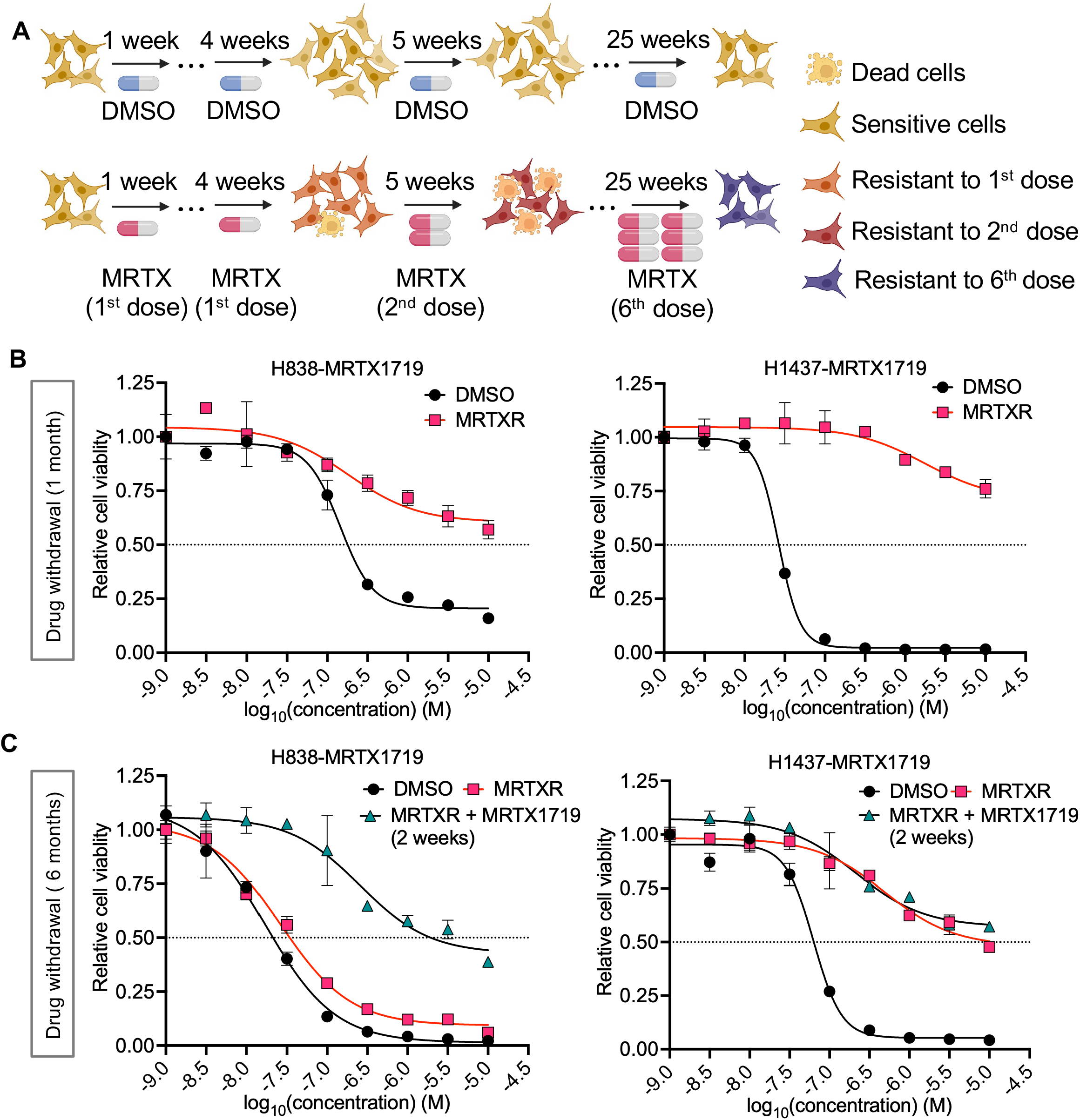
Acquired resistance to MRTX1719 in sensitive NSCLC cells *in vitro*. **(A)** Schematic illustrating the dose-escalation strategy used to generate MRTX1719-resistant cells. The concentration of MRTX1719 was doubled every 4 weeks. The starting concentrations for H838 and H1437 cells were 0.02 μM and 0.1 μM, respectively, and were escalated to final concentrations of 1 μM and 2 μM. **(B-C)** After 6 months of continuous MRTX1719 treatment, resistant cells were cultured without drug to assess the stability of resistance. MRTX1719 dose-response assays were performed in parental (DMSO) and MRTX1719-resistant (MRTXR) cells after 1 month of drug withdrawal **(B)**, and in DMSO, MRTXR, and MRTXR cells re-treated with MRTX1719 for 2 weeks after 6 months of drug withdrawal **(C).** Data are presented as mean ± SD.

During dose escalation, MRTX1719-treated cells became enlarged and exhibited markedly slower proliferation than DMSO-treated controls (data not shown). After long-term treatment, cells were cultured in drug-free medium for 1 month to allow growth recovery and were then tested for MRTX1719 sensitivity. Compared with DMSO-treated controls, MRTX1719-treated cells (MRTXR) showed clear acquired resistance in H838 and H1437 cells (**Figure 2B**), whereas A549 cells exhibited only a modest shift in sensitivity (data not shown). Given the limited change in A549 cells, this model was not investigated further. To assess the stability of this resistance, H838-MRTXR and H1437-MRTXR cells were maintained in drug-free medium and subjected to several freeze-thaw cycles. After 6 months of drug withdrawal, H838-MRTXR cells had lost most of their resistance, whereas H1437-MRTXR cells retained resistance, indicating a reversible resistant state in H838-MRTXR and a stable resistant state in H1437-MRTXR (**Figure 2C**). Following 2 weeks of MRTX1719 re-challenge, H838-MRTXR cells reacquired resistance, whereas H1437-MRTXR cells maintained a similar degree of resistance (**Figure 2C**). These results demonstrate that prolonged MRTX1719 treatment induces acquired resistance in sensitive NSCLC cells, with distinct reversible and stable resistance states.

### 3.3 MRTX1719 resistance is not fully explained by altered PRMT5 activity or reduced MTA levels

Having established MRTX1719-resistant models, we first examined whether acquired resistance was associated with altered PRMT5 activity. DMSO- and MRTXR-paired cells were treated with increasing doses of MRTX1719 for 6 days. SDMA reduction kinetics were broadly similar between sensitive and resistant cells, with a slight delay in SDMA loss in resistant cells at concentrations below 0.1 μM. Basal SDMA levels and PRMT5 expression were also largely unchanged in resistant cells (**Figure 3A**). Several bands were more intense in resistant cells at baseline, but this pattern was lost after extended culture in drug-free medium, suggesting that it was transient. Given the reported functional overlap between PRMT5 and PRMT1 (40-42), we also examined whether resistance was associated with compensatory changes in other PRMT activities. MMA and ADMA levels showed no obvious differences at baseline or after MRTX1719 treatment (**Figure 3A**), indicating that no major global compensatory changes in other PRMT activities were evident. We further assessed whether acquired PRMT5 mutations might impair MRTX1719 binding. Sanger sequencing of the PRMT5 coding sequence from both DMSO and MRTXR cells identified no coding mutations (data not shown). Together, these findings suggest that MRTX1719 resistance is not readily explained by overt changes in PRMT5 activity, compensatory changes in other PRMT activities, or PRMT5 coding mutations.

**Figure 3.**
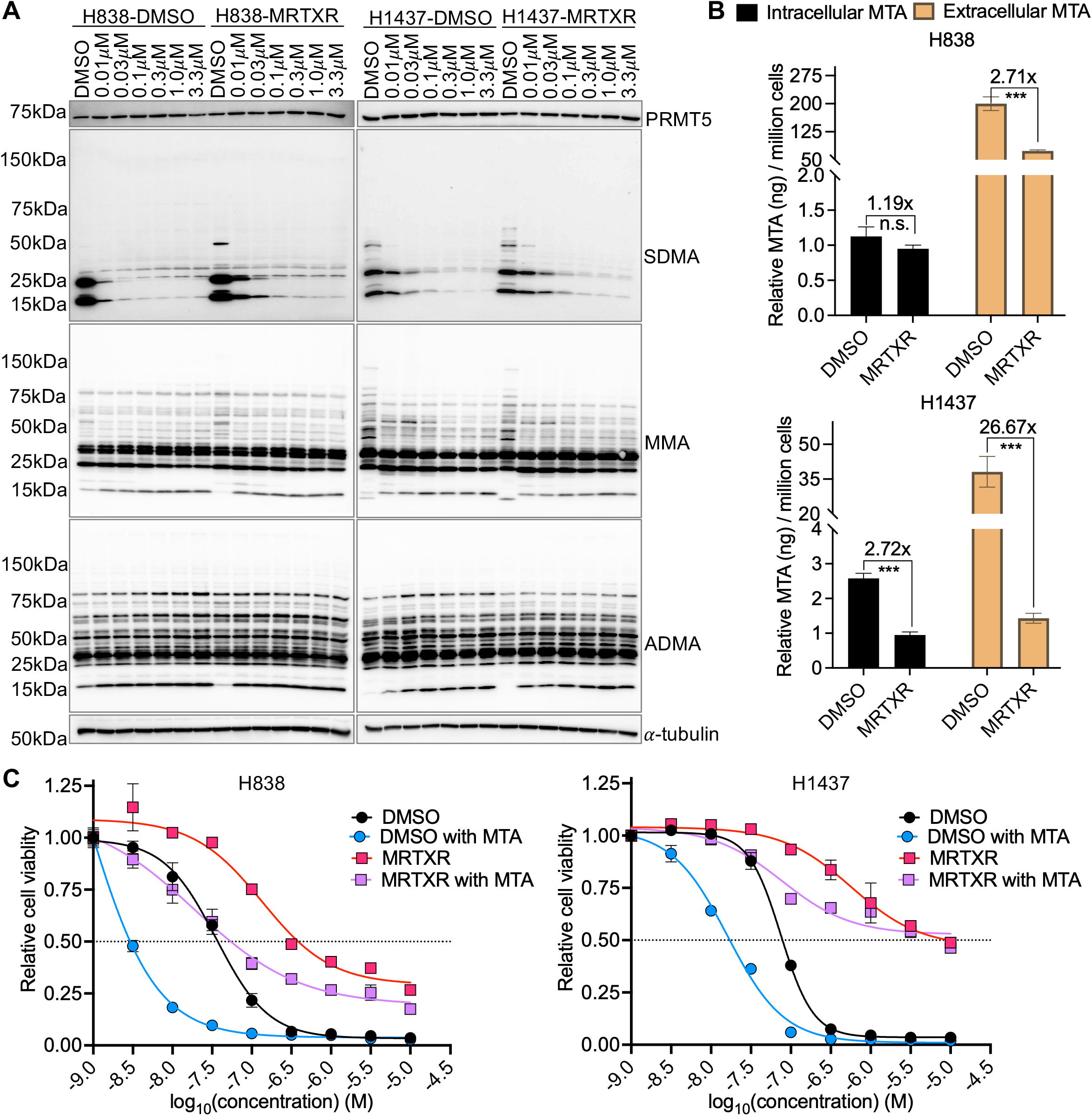
MRTX1719 resistance not fully explained by altered PRMT5 activity or reduced MTA levels. **(A)** Immunoblot analysis of PRMT5 and arginine methylation marks (SDMA, MMA, and ADMA) in DMSO and MRTXR cells treated with increasing concentrations of MRTX1719. **(B)** Quantification of intracellular and extracellular MTA levels in DMSO and MRTXR cells. **(C)** MRTX1719 dose-response assays in DMSO and MRTXR cells with or without MTA supplementation. Data are presented as mean ± SD. n.s., not significant; \**p* < 0.05; \*\**p* < 0.01; \*\*\**p* < 0.001.

We next examined whether altered MTA levels contribute to resistance. Intracellular and extracellular MTA levels were quantified by LC-MS in DMSO and MRTXR cell pellets and supernatants. Intracellular MTA was significantly decreased only in H1437-MRTXR cells, whereas extracellular MTA was significantly reduced in both resistant lines, with a more pronounced decrease in H1437-MRTXR (**Figure 3B**). To test whether reduced MTA drives resistance, DMSO- and MRTXR-paired cells were treated with MRTX1719 in the presence or absence of exogenous MTA and subjected to a 6-day viability assay. In DMSO cells, 10 μM MTA supplementation enhanced the growth-inhibitory effect of MRTX1719, consistent with its MTA-dependent mechanism of action. In resistant cells, 10 μM MTA supplementation partially restored sensitivity to MRTX1719, but did not fully reverse the resistant phenotype (**Figure 3C**). These results suggest that reduced MTA levels may contribute to, but are insufficient to explain, acquired resistance to MRTX1719.

### 3.4 Acquired MRTX1719 resistance confers collateral sensitivity to MEK inhibitors

As acquired resistance was not explained by altered PRMT5 activity or reduced MTA levels, we performed a high-throughput drug screen to identify therapeutic vulnerabilities associated with resistance. The screening library comprised 619 compounds, including SGC epigenetic compounds, TargetMol epigenetic inhibitors, and FDA-approved oncology drugs (**Figure S2A**). Screen performance was supported by the expected responses to the positive control MRTX1719 and the nonselective control anisomycin (**Figure S2B**).

Differential drug response analysis identified compounds to which H838- and H1437-MRTXR cells were either more or less sensitive than their corresponding controls (**Figure 4A, Supplementary Tables 2 and 3**). Five significantly sensitizing hits overlapped between the two resistant lines, including four MEK inhibitors and the mTOR inhibitor rapamycin. By contrast, one overlapping hit associated with reduced sensitivity, the SAM-competitive PRMT5 inhibitor LLY-283, was identified in both resistant lines, indicating cross-resistance among distinct classes of PRMT5 inhibitors (**Figure 4B**). We next validated this collateral sensitivity to MEK inhibition in DMSO- and MRTXR-paired cells. Dose-response analysis showed that H838- and H1437-MRTXR cells were more sensitive than control cells to trametinib (**Figure 4C**) and selumetinib (**Figure S2C**), consistent with the screening results.

**Figure 4.**
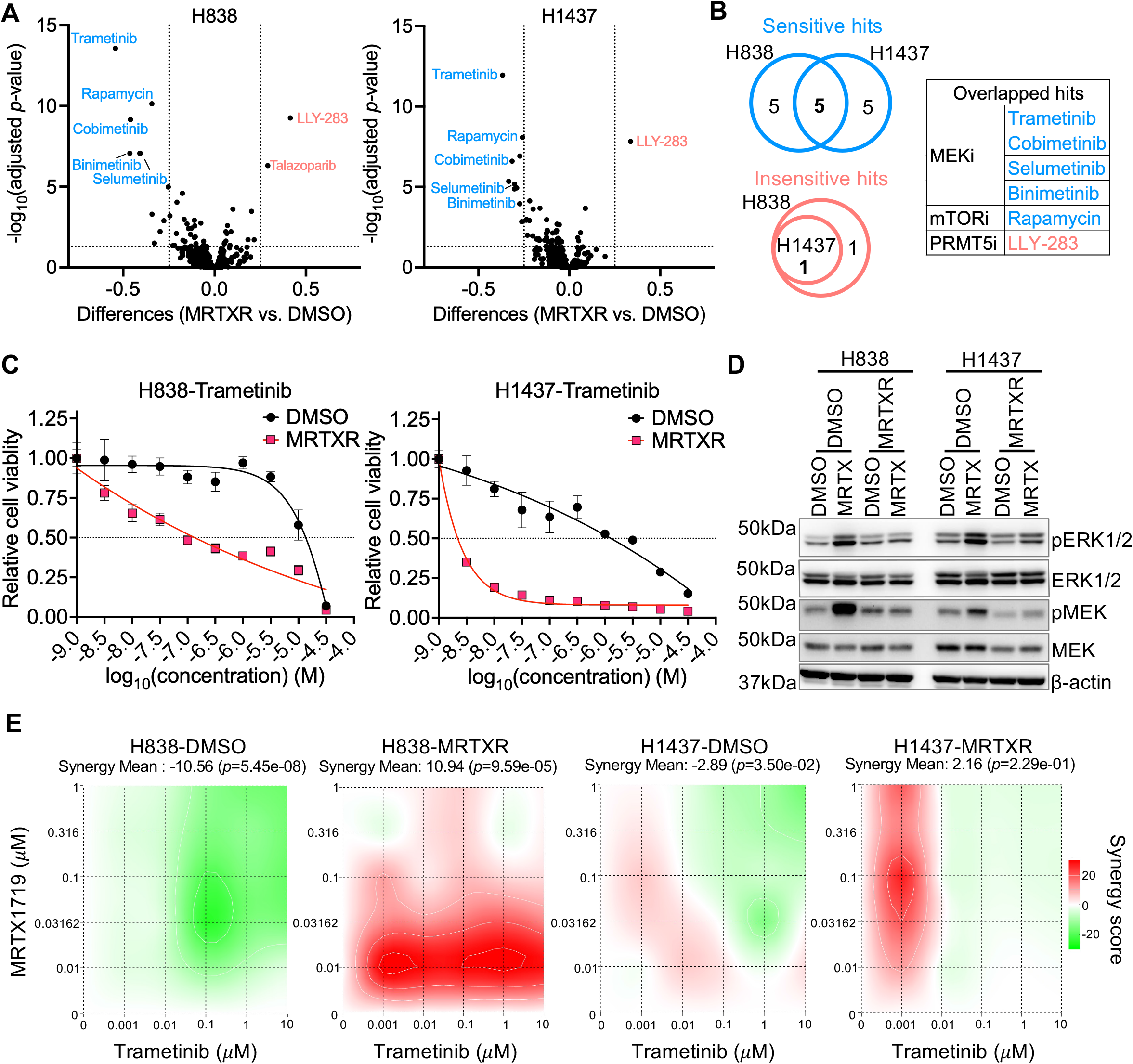
Collateral sensitivity to MEK inhibitors in MRTX1719-resistant NSCLC cells *in vitro*. **(A)** Volcano plots illustrating differential drug sensitivity between MRTXR and DMSO cells from the high-throughput drug screen. Significant hits were defined as viability differences (MRTXR vs. DMSO) > 0.25 or < −0.25 with an adjusted *p*-value < 0.05. Blue dots indicate drugs to which MRTXR cells are more sensitive than DMSO cells, whereas red dots indicate drugs to which DMSO cells are more sensitive. **(B)** Venn diagram (left) and summary table (right) showing overlapping significant hits identified in both cell lines. **(C)** Dose-response curves of DMSO and MRTXR cells treated with the MEK inhibitor trametinib. Data are presented as mean ± SD. **(D)** Immunoblot analysis of total and phosphorylated ERK and MEK in DMSO and MRTXR cells with or without MRTX1719 treatment for 3 days. **(E)** Synergy analysis of MRTX1719 and trametinib in DMSO and MRTXR cells. Synergy mean scores were calculated using the Bliss model with the SynergyFinder+ tool.

We next examined MAPK signaling by assessing MEK and ERK activity. Basal MEK and ERK phosphorylation were not obviously increased in resistant cells relative to sensitive controls. However, after 3 days of MRTX1719 treatment, MEK and ERK phosphorylation increased in H838 and H1437 sensitive cells, whereas this response was blunted in resistant cells (**Figure 4D**). These findings suggest that acute MRTX1719 treatment activates MAPK signaling in sensitive cells, while prolonged treatment is associated with an altered MAPK response in resistant cells.

We then tested the effect of combining MRTX1719 with MEK inhibitors trametinib and selumetinib on DMSO- and MRTXR-paired cells. In sensitive cells, synergy mean scores indicated an antagonistic trend rather than synergy. By contrast, resistant cells displayed synergistic responses to combined MRTX1719 and MEK inhibition, particularly in H838-MRTXR cells and, at low doses, in H1437-MRTXR cells (**Figure 4E, S2D**). Together, these findings identify collateral sensitivity to MEK inhibition as a therapeutic vulnerability of acquired MRTX1719 resistance.

### 3.5 MAPK-related transcriptional programs are enriched in MRTX1719-resistant cells

To characterize global transcriptional changes associated with acquired MRTX1719 resistance, we performed RNA sequencing in DMSO- and MRTXR-paired cells. Using |log_2_ fold change| > 1 and FDR < 0.05 to define significantly differentially expressed coding genes in MRTXR relative to DMSO cells, we identified 574 upregulated genes in H838 and 937 in H1437, with 74 genes shared between the two models. Similarly, 558 downregulated genes in H838 and 1,134 in H1437 were identified, with an overlap of 111 genes (**Figure 5A, Supplementary Tables 4 and 5**). Fisher’s exact test indicated that the overlaps between the two cell models were significant for both upregulated (*p*-value = 2.02 x 10^-9^) and downregulated genes (*p*-value = 8.39 x 10^-22^).

**Figure 5.**
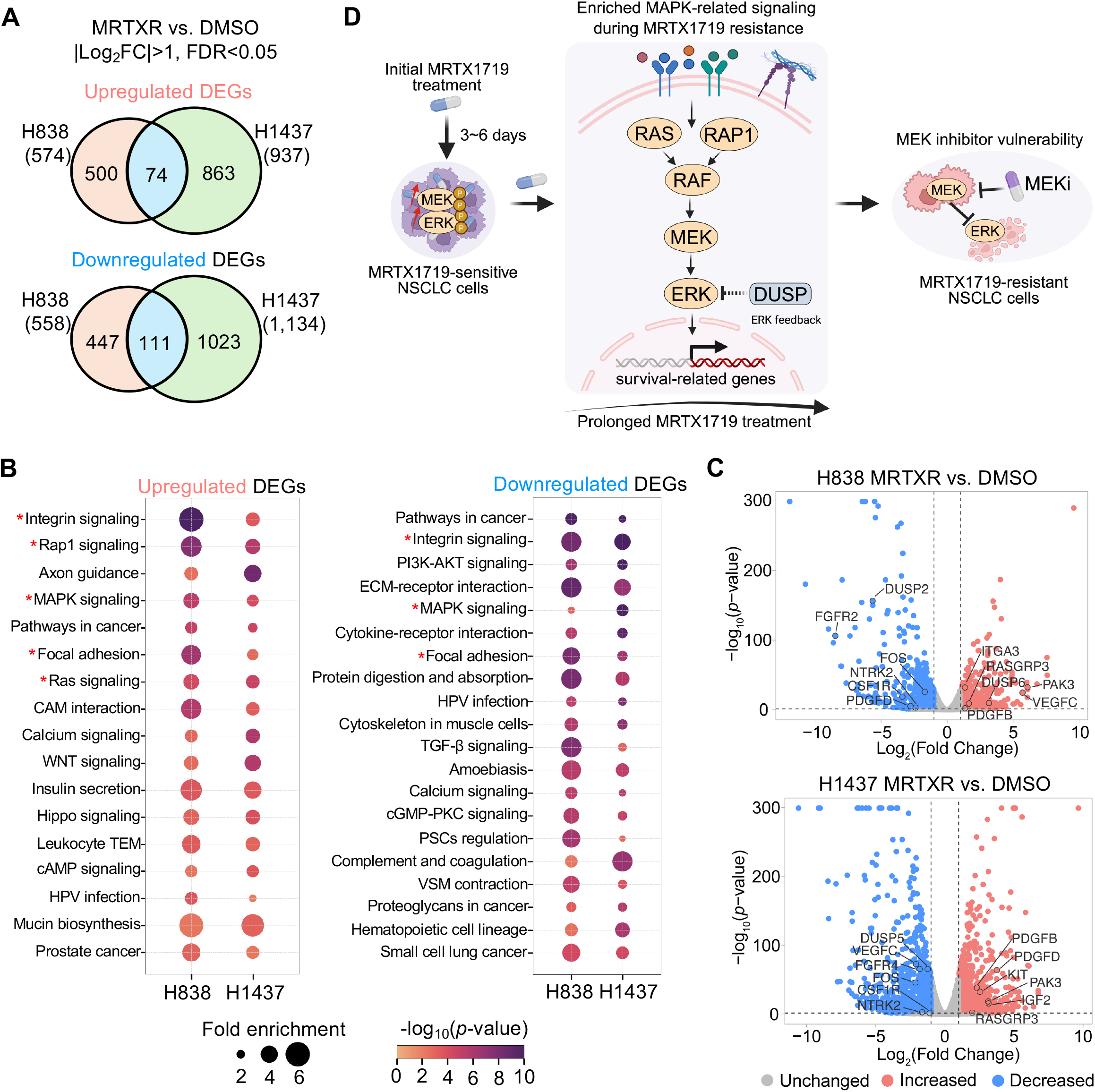
Enriched MAPK-related transcriptional programs in MRTX1719-resistant cells. **(A)** Venn diagram showing the number of overlapping differentially expressed genes (DEGs) identified by RNA-seq analysis in both cell lines. DEGs were defined as coding genes with |log_2_ (fold change, FC)| > 1 between MRTXR and DMSO cells with FDR < 0.05. Fisher’s exact test was used to assess the significance of the overlap between the two cell models for upregulated genes (*p-value* = 2.02 x 10^-9^) and downregulated genes (*p*-value = 8.39 x 10^-22^). **(B)** KEGG pathway enrichment analysis of upregulated (left) and downregulated (right) DEGs identified in each cell line. The top 20 enriched pathways are shown for downregulated DEGs. MAPK-related pathways are marked with asterisks. **(C)** Representative DEGs in MAPK-related transcriptional programs shown in volcano plots. **(D)** Proposed model illustrating that MRTX1719 initially activates MAPK signaling, and continued long-term treatment is associated with the emergence of a resistant state characterized by enrichment of MAPK-related transcriptional programs and collateral sensitivity to MEK inhibition.

We next performed KEGG pathway enrichment analysis using Metascape (37) on the upregulated and downregulated gene sets from each cell line. MAPK-related pathways, including MAPK signaling itself as well as upstream or associated pathways such as RAS, RAP1, integrin, and extracellular adhesion signaling, were significantly enriched in both resistant models (**Figure 5B**). Representative differentially expressed genes in these pathways were shown in the volcano plots (**Figure 5C**). Notably, both positive and negative regulators of MAPK-related pathways were differentially expressed in resistant cells. These findings suggest that MAPK signaling is not simply increased, but rather transcriptionally altered, in resistant cells.

Together, these data show that MRTX1719-resistant cells exhibit enrichment of MAPK-related transcriptional programs, consistent with the collateral sensitivity to MEK inhibition identified in the drug screen. Based on these observations, we propose a working model in which MRTX1719 initially activates MAPK signaling, and continued long-term treatment is associated with the emergence of a resistant state characterized by enrichment of MAPK-related transcriptional programs and collateral sensitivity to MEK inhibition (**Figure 5D**).

## 4. Discussion

In this study, we generated multiple MTAP-isogenic cell models to evaluate the contribution of MTAP loss to MRTX1719 efficacy. Although MRTX1719 showed preferential activity in MTAP-KO cells relative to MTAP-WT counterparts, responses among MTAP-KO models were variable. Importantly, these MTAP-isogenic models provide a valuable resource for demonstrating that MTAP loss alone is insufficient to predict response to MRTX1719, consistent with the modest ORRs observed clinically (43). Future studies should therefore focus on identifying predictive biomarkers and rational combination strategies to improve the efficacy of MRTX1719 more broadly. In this context, several combinations of MRTX1719 with chemotherapy, targeted therapy and immunotherapy are currently under clinical evaluations (43).

To investigate acquired resistance to MRTX1719 and identify associated vulnerabilities, we established resistant cell models through prolonged drug exposure. These models displayed distinct resistance phenotypes. H838 cells showed a reversible resistant state, losing resistance after prolonged drug withdrawal and regaining it upon re-treatment, whereas H1437 cells retained resistance throughout withdrawal, consistent with a stable resistant state. These findings suggest that resistance to MRTX1719 may arise through distinct, cell context-dependent mechanisms, potentially involving reversible adaptive plasticity in H838 cells and more durable molecular alterations in H1437 cells, as reported for other targeted therapies (44-50). Because our mechanistic analysis was limited to the PRMT5 coding sequence, future studies integrating whole-genome, transcriptomic, and epigenetic analyses will be needed to define additional genetic and non-genetic drivers of resistance.

A key finding of this study is that acquired resistance to MRTX1719 confers collateral sensitivity to MEK inhibitors. This vulnerability was supported by RNA-sequencing data showing enrichment of MAPK-related transcriptional programs in resistant cells. However, basal phosphorylated MEK and ERK levels were not obviously elevated in resistant cells compared with paired control cells. Instead, acute MRTX1719 treatment induced MEK and ERK phosphorylation, whereas this response was blunted in resistant cells. These results suggest that prolonged MRTX1719 exposure does not simply cause sustained MAPK activation, but instead leads to adaptive remodeling of MAPK signaling that creates vulnerability to MEK inhibition.

The mechanism linking PRMT5 inhibition to MAPK-related signaling remains unclear. One possibility is a direct effect on MAPK pathway components, as PRMT5-dependent methylation has been reported to promote the degradation of activated CRAF and BRAF (51). Another possibility is indirect regulation through pathway crosstalk. Notably, downregulated genes in resistant cells were enriched in the PI3K-AKT pathway. This is of interest because PRMT5-mediated arginine methylation has been shown to activate AKT kinase and promote tumorigenesis and metastasis (52,53). Further investigation of RAF and AKT activity will be needed to clarify how PRMT5 inhibition remodels signaling in resistant cells.

Based on the induction of MAPK signaling following short-term MRTX1719 treatment in sensitive cells, we evaluated the combination of MRTX1719 and MEK inhibition. Unexpectedly, this combination showed an antagonistic trend in sensitive cells, whereas resistant cells exhibited synergy in H838-MRTXR cells and, at low doses, in H1437-MRTXR cells. Previous studies reported synergistic effects of MRTX1719 combined with trametinib in KRAS-mutant NSCLC models and of MRTX1719 combined with KRAS inhibition in KRAS-mutant PDAC models (54,55). Our findings are not necessarily inconsistent with these reports, but may reflect differences in genetic and signaling context. Unlike those KRAS-mutant models, H838 and H1437 cells are KRAS- and RAF-wildtype. In KRAS-mutant settings, constitutive MAPK activation may underlie the observed synergy, whereas in our models MAPK dependency appears to emerge during acquisition of resistance. These results raise the possibility that sequential treatment with MRTX1719 followed by MEK inhibition may be a more effective strategy in KRAS-wildtype tumors, although this will require validations in *in vivo*.

## Supplementary Materials

Detailed information on the generated MTAP-isogenic cells, the drug screening results and RNA sequencing results is provided in Supplementary Tables 1-5.

## Data Availability

The data generated in this study are available upon request from the corresponding author.

## Acknowledgments

This work was supported by grants from the National Institutes of Health (H.X., 1R35GM137927; J.R. & H.X., 1R21CA296234), Department of Defense (T.C., W.P. & H.X., ME220215), American Cancer Society (W.P., RSG-23-1155993-01-MM), CPRIT Core Facility Support Awards (C.S., RP200668 and RP250505) and MD Anderson Cancer Center Support Grant (J.R., M.B. & H.X., P30 CA016672). H.X. is a CPRIT Scholar in Cancer Research.

## Author Contributions

H.X., M.B. and R.F. conceived the study. R.F. and Y.W. performed experiments, analyzed and interpreted the data, and prepared the original draft. E.B., I.R. and R.F. generated and characterized the MTAP-isogenic cell lines. S.S. and X.L. performed LC-MS experiments and analyzed the data. N.N., R.P. and C.S. performed the high-throughput drug screening and analyzed the data. A.A. and W.L. repeated drug sensitivity assays and confirmed the resistant phenotype. S.W., W.H., Y.L., and B.L. processed and analyzed the RNA-seq data. P.S., J.R., T.C. and W.P. contributed conceptual input. ChatGPT was used solely for minor editing to correct typographical and grammatical errors.

## Figure Legends

**Figure S1.**
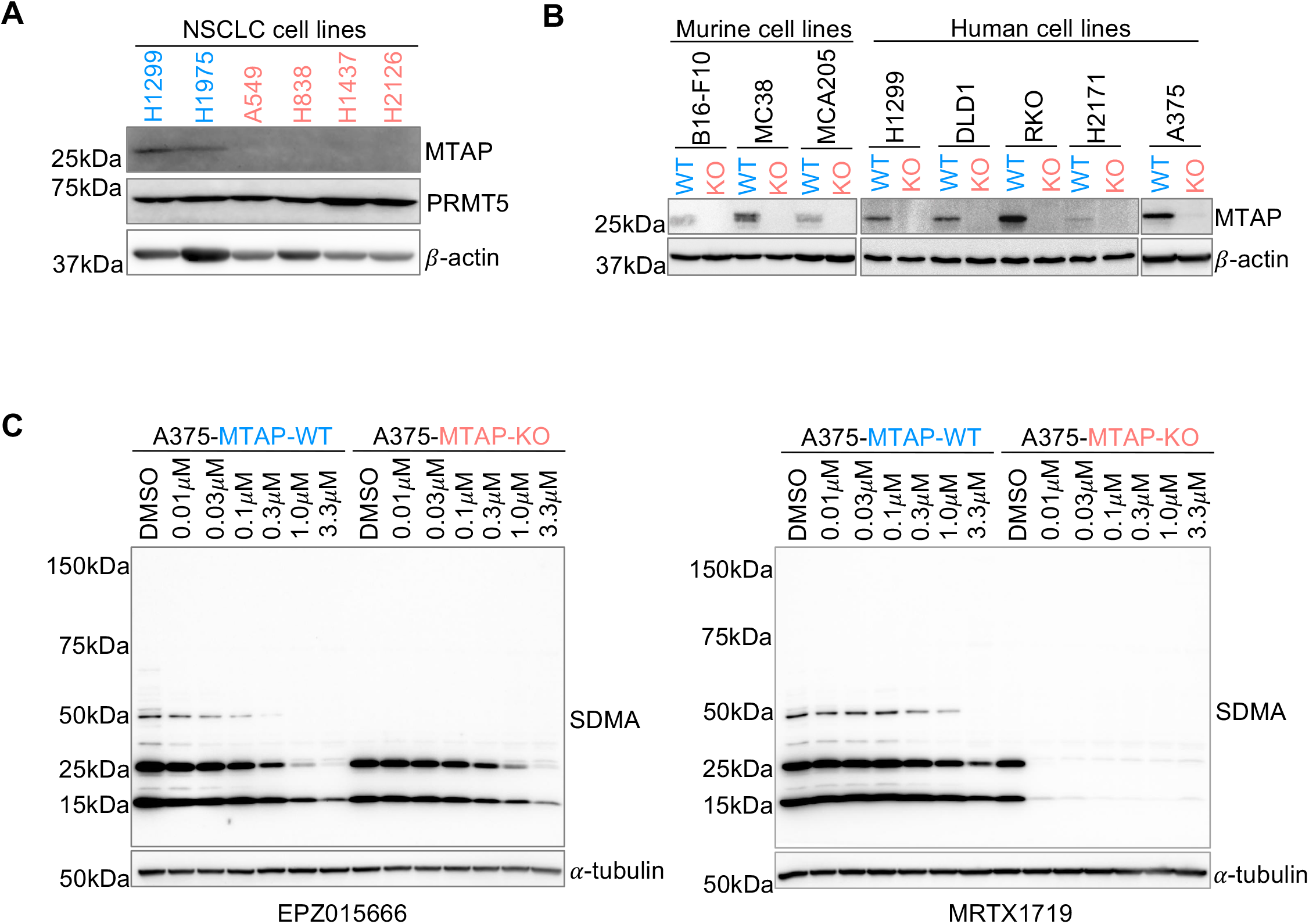
Assessment of MTAP status and PRMT5 inhibition in MTAP-isogenic cells. **(A)** Immunoblot analysis of MTAP and PRMT5 protein levels across a panel of NSCLC cell lines. **(B)** Immunoblot validation of MTAP expression in MTAP-isogenic murine and human cell lines. **(C)** Immunoblot analysis of SDMA levels in the A375 MTAP-isogenic cell pair following treatment with increasing concentrations of EPZ015666 or MRTX1719.

**Figure S2.**
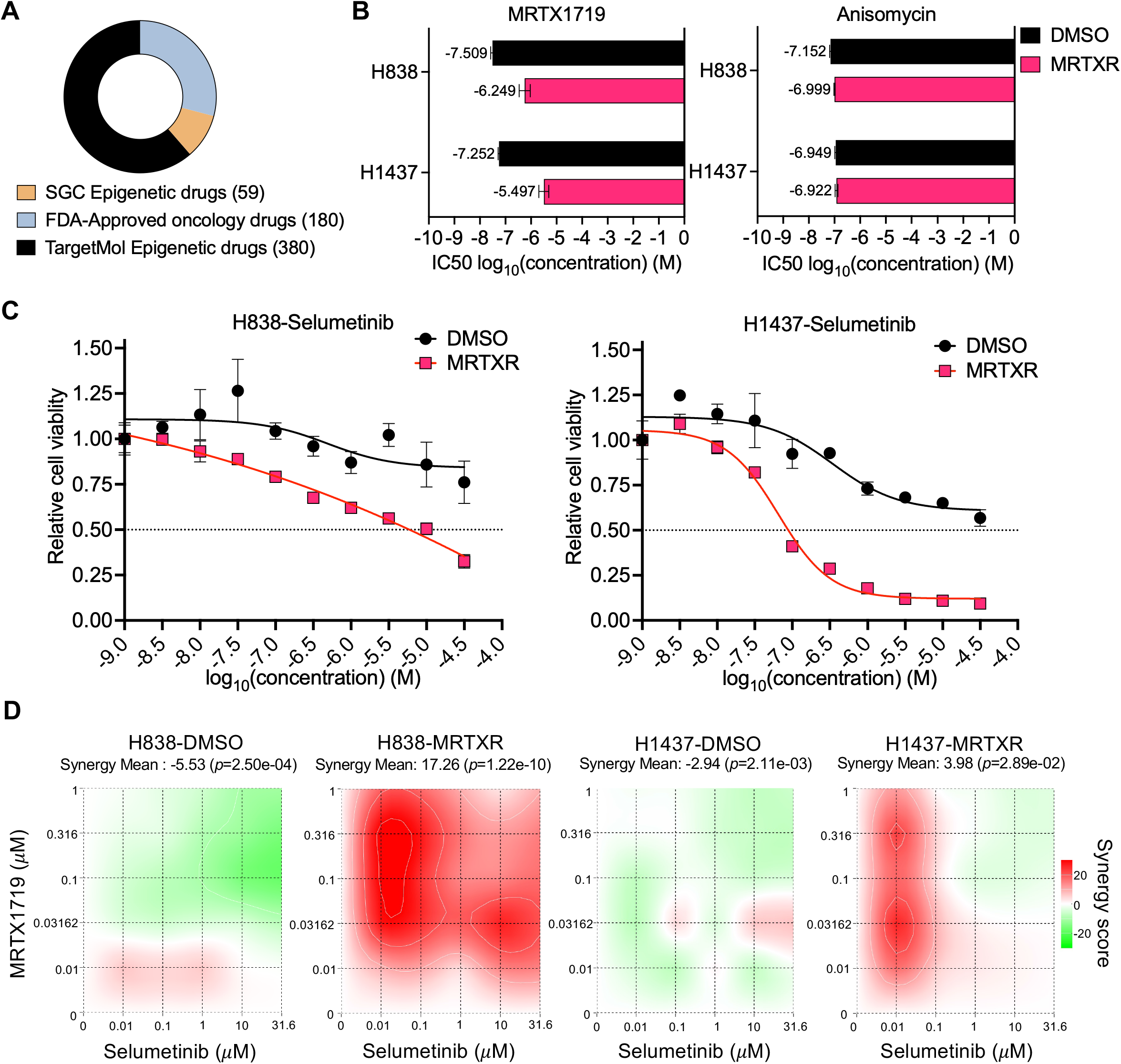
High-throughput drug screen and MEK inhibitor sensitivity in MRTX1719-resistant NSCLC cells. **(A)** Composition of the compound library used for drug screen, including SGC epigenetic compounds, FDA-approved oncology drugs, and TargetMol epigenetic inhibitors. **(B)** IC50 values of MRTX1719 and anisomycin in DMSO and MRTXR cells. Anisomycin was included as a nonselective control in the drug screen. **(C)** Dose-response curves of DMSO and MRTXR cells treated with the MEK inhibitor selumetinib. Data are presented as mean ± SD. **(D)** Synergy heatmaps of MRTX1719 and selumetinib in DMSO and MRTXR cells. Synergy mean scores were calculated using the Bliss model with the SynergyFinder+ tool.

